# Acute vagus nerve stimulation does not affect liking or wanting ratings of food in healthy participants

**DOI:** 10.1101/2021.03.26.437062

**Authors:** Franziska K. Müller, Vanessa Teckentrup, Anne Kühnel, Magdalena Ferstl, Nils B. Kroemer

## Abstract

The vagus nerve plays a vital role in the regulation of food intake and vagal afferent signals may help regulate food cue reactivity by providing negative homeostatic feedback. Despite strong evidence from preclinical studies on vagal afferent “satiety” signals in guiding food intake, evidence from human studies is largely inconclusive to date. Here, we investigated the acute effects of left or right transcutaneous auricular vagus nerve stimulation (taVNS) on subjective ratings of wanting and liking of various food and non-food items in 82 healthy participants (46 women, M_BMI_ = 23.1 kg/m^2^). In contrast to previous reports in patients with depression, we found moderate to anecdotal evidence supporting the absence of taVNS-induced changes in food ratings. To test whether the absence of taVNS effects on food ratings is due to heterogeneity in the sample, we conducted post hoc subgroup analyses by splitting the data according to stimulation side and sex (between-subject factors) as well as caloric density, perceived healthiness, and flavor (sweet vs. savory) of the food (within-subject factors). This multiverse analysis largely supported the absence of taVNS-induced changes since the strongest subgroup effects provided only anecdotal evidence in favor of taVNS-induced changes. We conclude that acute taVNS only has a marginal effect on subjective ratings of food, suggesting that it is an unlikely mechanism for the reported long-term effects of VNS on body weight. In light of an absence of acute taVNS effects on food craving, our results call for future research on the correspondence between acute and chronic effects of vagal afferent stimulation.

## Introduction

Excessive food intake over prolonged periods of time is associated with metabolic disorders such as obesity which has become more prevalent in modern times (GBD Obesity Collaborators et al., 2017; David Val-Laillet et al., 2015). Most attempts to lose weight are not successful because it is hard to resist food that we like (de Araujo, Schatzker, & Small, 2019; Lowe et al., 2009). Intriguingly, the vagus nerve has been shown to play an essential role in the regulation of food intake by facilitating gut-brain interactions (Berthoud, 2008; Breit, Kupferberg, Rogler, & Hasler, 2018; Cork, 2018; de Araujo, Ferreira, Tellez, Ren, & Yeckel, 2012; Guillaume de Lartigue, 2016). Nevertheless, it is largely elusive how vagal afferent signals are involved in food craving in humans. Therefore, studying ratings of food cue-induced wanting or liking during transcutaneous auricular vagus nerve stimulation (taVNS) is promising for understanding the contribution of vagal afferent signals to food craving.

The vagus nerve is the tenth and longest cranial nerve, arising from the brainstem and spreading through the head, neck, chest and abdomen (Breit et al., 2018; Yuan & Silberstein, 2016). Afferent vagal signaling provides information about meal sizes and digested nutrients to appetite and satiety centers in the brain (Berthoud, 2008; Bonaz, Picq, Sinniger, Mayol, & Clarençon, 2013; Breit et al., 2018; Browning, Verheijden, & Boeckxstaens, 2017; Kaniusas et al., 2019). Vagal anorexigenic fibers are mechanosensitive and chemosensitive (Wang, de Lartigue, & Page, 2020), forwarding signals elicited by stomach distension and digested nutrients. This feedback on food intake is conveyed to the brain and leads to an inhibition of consumption (Guillaume de Lartigue, 2016). Remarkably, chronic high-caloric diet can result in a disturbance of such vagal feedback mechanisms which might contribute to overeating and obesity (Guillaume de Lartigue, 2016). In addition to chemical and mechanical signals from the gut, vagal afferent projections are also affected by metabolic hormones such as ghrelin or cholecystochinine (Guillaume de Lartigue, 2016; Morton, Cummings, Baskin, Barsh, & Schwartz, 2006; Palmiter, 2007). Furthermore, efferent vagal signaling regulates digestion (Breit et al., 2018). Based on an afferent-efferent feedback loop, this provides a way for afferent signals to alter energy metabolism (Teckentrup et al., 2020). Collectively, these studies highlight the vital contribution of the vagus nerve in gut-brain signaling to guide food intake according to homeostatic needs.

In line with the vital role of vagal afferent signals in regulating eating behavior, vagus nerve stimulation (VNS) has been conclusively shown to affect food intake and induce weight loss in animals (Bugajski et al., 2007; Gil, Bugajski, & Thor, 2011; Li et al., 2015; Roslin & Kurian, 2001; Sobocki, Krolczyk, Herman, Matyja, & Thor, 2005; D. Val-Laillet, Biraben, Randuineau, & Malbert, 2010; Yao et al., 2018). In humans, however, results are largely inconsistent (Pelot & Grill, 2018). One study reported altered craving for sweet foods due to acute cervical VNS in depressed patients (Bodenlos et al., 2007). A small pilot study in healthy participants found that taVNS increased liking of low-fat, but not high-fat puddings, compared to sham stimulation (Öztürk, Büning, Frangos, De Lartigue, & Veldhuizen, 2020). Notably, other studies observed no effects on food consumption or ratings (Alicart et al., 2020; Obst, Heldmann, Alicart, Tittgemeyer, & Munte, 2020). Beyond food intake, vagal afferent signals can alter motivational behavior more broadly due to effects on monoaminergic neurotransmission including mesolimbic dopamine (de Araujo et al., 2012; Han et al., 2018). Consequently, we have recently shown that acute taVNS alters reward-related learning (Kühnel et al., 2020) and boosts the drive to work for food and monetary rewards (Neuser et al., 2020). Although these findings may suggest corresponding changes in ratings, we did not observe an increase in subjective wanting, but a greater willingness to work for rewards that were less wanted (Neuser et al., 2020). Thus, there is growing albeit inconclusive evidence from preclinical work suggesting that VNS as well as taVNS could alter food craving.

To summarize, vagal signaling has been repeatedly shown to regulate eating behavior. However, the acute impact of vagal afferent stimulation on food intake in humans is still largely inconclusive (Pelot & Grill, 2018) and the mechanisms leading to weight loss after chronic VNS remain elusive (Abubakr & Wambacq, 2008; Burneo, Faught, Knowlton, Morawetz, & Kuzniecky, 2002; Pardo et al., 2007). To close this gap, we examined the impact of acute taVNS on cue-induced food craving in healthy participants. In contrast to previous studies primarily focusing on invasive cervical VNS, we non-invasively stimulated the auricular branch of the vagus nerve at the cymba conchae of the external ear (Ellrich, 2011; Peuker & Filler, 2002). To measure the effects of taVNS on food craving, we used a food-cue reactivity (FCR) task that included ratings of subjective liking and wanting. We hypothesized that taVNS might reduce cue-induced craving for food since vagal afferent signals contribute to a physiological feedback loop terminating intake after a meal (G. de Lartigue, 2016).

More specifically, we expected taVNS compared to sham stimulation to reduce subjective ratings of wanting as taVNS has been shown to reduce craving for sweet food before (Bodenlos et al., 2007). In contrast, we did not expect taVNS-induced changes in anticipatory liking ratings due to the absence of post-ingestive feedback signals in our design (Öztürk et al., 2020). To assess the specificity for food cues of potential taVNS effects, wanting and liking ratings for non-food items served as a control condition. In line with our hypothesis, ratings of liking were not affected by acute taVNS. Contrary to previous results, however, acute taVNS did not affect subjective ratings of wanting, which is in line with previous results derived from an effort allocation task within the same study (Neuser et al., 2020). The absence of taVNS effects was corroborated by Bayesian analyses including potential modulating factors such as stimulation side and sex (between-subject factors) and caloric density, perceived healthiness, and flavor (sweet vs. savory) of the food (within-subject factors).

### Methods

#### Participants

For the current study, we enrolled 85 healthy participants who had been recruited via the university’s mailing list and flyers (for details, see Neuser et al., 2020). The study was approved by the local ethics committee according to the Declaration of Helsinki and each participant gave written informed consent. Participants were included if they were between 18 and 40 years old, right-handed, and German speaking. Participants were excluded if they suffered from diabetes, severe brain injuries, schizophrenia, bipolar disorder, major depressive disorder, a moderate or severe substance use disorder as well as any anxiety disorder (except specific phobia), obsessive compulsive disorder, posttraumatic stress disorder, somatic symptom disorder or any eating disorder within the past 12 months. To verify the participant’s eligibility for the study, we conducted screenings by phone. The participants were compensated with either €32 or partial course credits plus additional money and breakfast as well as snacks depending on their performance in two other tasks that followed the FCR task. For the reported analyses, we excluded three participants because they did not complete the second session. Consequently, we analyzed data from 82 participants (46 women, M_age_ = 24.4 years, M_BMI_ = 23.1 kg/m^2^, [17.9, 30.9], M_WtHR_= 0.45, [0.36, 0.56]). Out of this sample, 42 participants received taVNS at the left ear, whereas 40 participants received taVNS at the right ear.

#### Experimental procedure

Each participant completed two sessions approximately at the same time in the morning after an overnight fast with a break of at least two and up to seven days in between. In one of the sessions, participants received taVNS, whereas they received sham stimulation in the other session. The order of the stimulation conditions was randomized. During the sessions, participants could drink as much water as they would like.

Each session included anthropometric measurements such as weight, height, hip and waist circumference as well as pulse, state questionnaires (Ferstl et al., 2020), and three computerized tasks. The first task was the FCR task in which participants rated their wanting and liking after viewing food pictures. The FCR task also included images of objects (i.e., stationery) as non-food control condition. Specific methods and results of the two other tasks (motivation: (Neuser et al., 2020); reinforcement learning: (Kühnel et al., 2020)) have been reported before. After the tasks, there was a 10 min break when participants received their breakfast (cereal) based on earned “energy points” in the second task. By the end of the session, participants received their compensation (money and/or partial course credit).

#### Transcutaneous vagus nerve stimulation device

To stimulate the auricular branch of the vagus nerve, we used Cerbomed NEMOS (Cerbomed GmbH, Erlangen) which delivers a 25 Hz stimulation in a biphasic 30 s on, 30 s off duty cycle. TaVNS was administered with the electrodes placed at the cymba conchae of the external ear which is strongly innervated by the auricular branch of the vagus nerve. For sham stimulation, the device was placed upside down so that the electrodes were positioned at the ear lobe (Frangos, Ellrich, & Komisaruk, 2015) which is not innervated by the vagus nerve (Peuker & Filler, 2002). To ensure adequate skin contact, we rubbed the skin with alcohol and placed medical tape to secure the electrodes in place.

For both conditions, the stimulation strength was individually adjusted. To this end, we used a pain visual analogue scale to track the participant’s sensations during the stepwise increase of the stimulation strength (steps of 0.1 or 0.2 mA) starting at 0.1 mA. The individual intensity for each participant then corresponded to a “tingling” sensation below the pain threshold (Frangos et al., 2015).

#### Food-cue task including ratings of liking and wanting

To measure food craving, we used a FCR task (Figure 1). Participants viewed pictures of food and stationery non-food control pictures and rated their liking and wanting of these items. We used a previously reported set of pictures (80 food items, 40 stationery items) provided by Charbonnier, van Meer, van der Laan, Viergever, and Smeets (2016). From this set of pictures, 60 food and 20 non-food pictures were randomly drawn for each participant and presented twice per session to measure liking or wanting, respectively. To avoid systematic order confounds, we randomized the order of the ratings and images. Due to a minor error in the randomization script, the ratings were not fully balanced (one wanting, one liking rating per image) for all participants. However, since we are primarily interested in the contrast between taVNS and sham sessions, this error does not systematically affect the comparisons between sessions.

**Figure 1:**
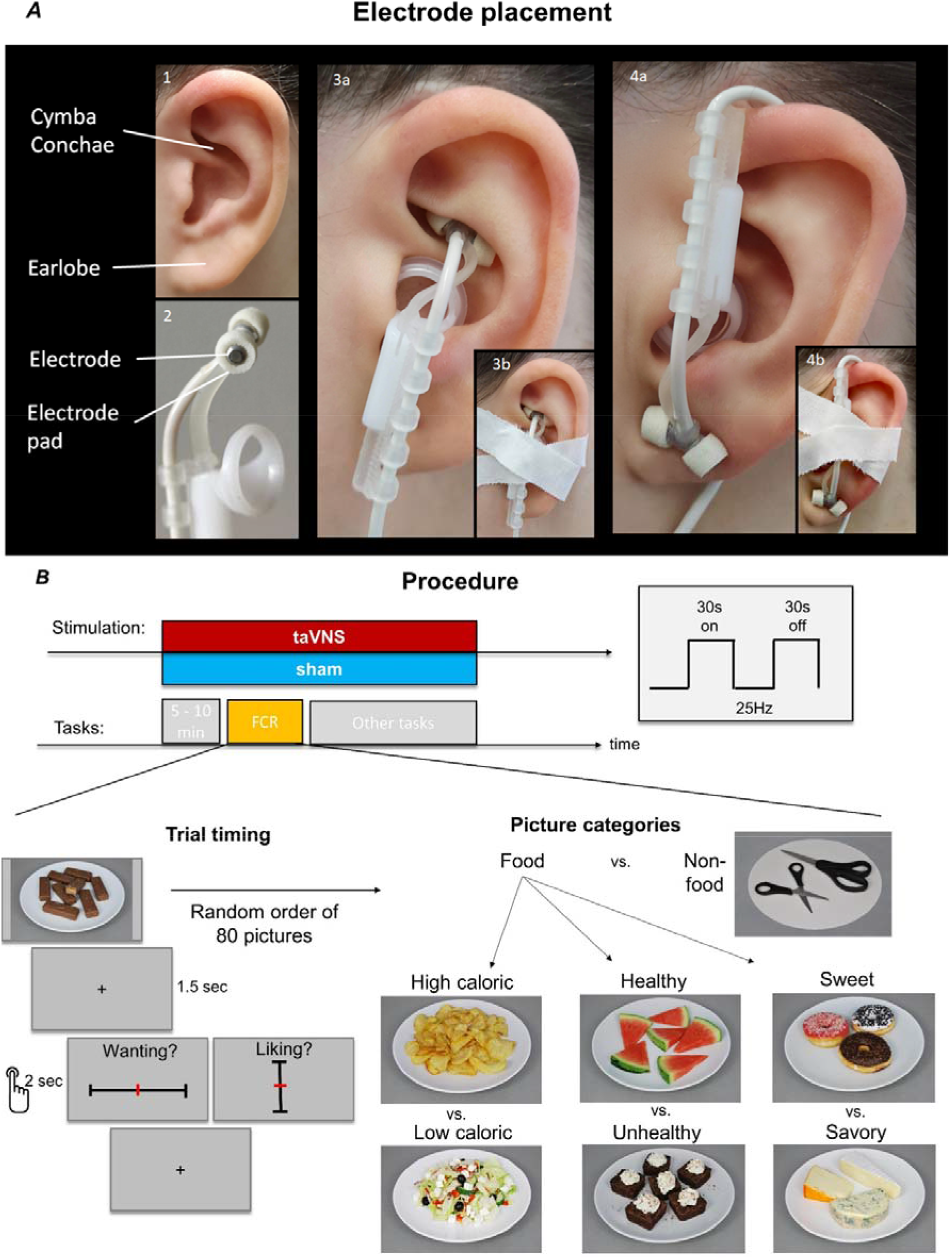
Summary of the experimental design. A: Electrode placement for the two conditions (taVNS and sham). B: Experimental procedure and duty cycle. As part of the experimental sessions, participants completed a food cue reactivity (FCR) task, where they viewed pictures of various food and non-food items. Examples for the categories are provided that were used for further subgroup analyses. After participants viewed the pictures, they rated their liking and wanting on vertical and horizontal visual analog scales, respectively.

The FCR task started approximately 5 to 10 minutes after stimulation onset. The images were shown for 1.5 s each. Then, participants viewed a black fixation cross on a white screen (inter-stimulus interval) before the rating scale appeared on the screen for a maximum of 2 s. After the rating was submitted, another fixation cross was shown (inter-trial interval). To keep the rate of stimuli the same, the intertrial interval was extended by the time that participants saved by pressing the button before the allotted 2 s had passed. Ratings were submitted by moving the left thumb joystick on an Xbox 360 controller. If participants did not submit the rating by pressing the (right thumb) A-button on the controller within 2 s, the last position on the scale was saved as rating. However, if participants did not move the joystick at all, such an unsubmitted rating was considered as invalid and removed from the analysis.

To measure liking ratings, we used a vertically labeled hedonic (visual analogue) scale. Liking ratings ranged from −100 (strongest disliking imaginable) to +100 (strongest liking imaginable; Lim, Wood, & Green, 2009). The participants were asked to “rate in the context of the full range of sensations that they have experienced in their life” and were provided with five gradual anchors on the scale’s axis from which the two extremes were labeled. Wanting ratings were acquired using a horizontal scale and ranged from 0 (not wanted at all) to 100 (strongly wanted). The different orientation (horizontal vs. vertical) and the delay between the ratings was intended to emphasize the differences between the two concepts.

#### Classification of food pictures

To analyze potential taVNS effects on specific kinds of food such as sweets, we used several classifications. First, we classified images according to caloric density of the depicted food as provided by Charbonnier et al. (2016). For one picture (#50, Grissini bread sticks), the caloric density was not provided by the authors, so we calculated 418kcal/100g based on similar items. Second, we used perceived healthiness on a 1-to-9-point rating scale collected in adult samples of four different countries as reported by Charbonnier et al. (2016). Third, we used the common categorization of sweet versus savory food to evaluate whether taVNS primarily affects linking or wanting of sweet foods.

#### Data analysis

To assess taVNS-induced changes in food craving, we ran one-sample t-tests on individual differences in average ratings between taVNS and sham sessions using both frequentist and Bayesian inference (i.e., equivalent to a paired t-test). As a measure of effect size, we used *d_z_* for within-subject effects. We used Bayesian inference to estimate the likelihood of the alternative hypothesis (taVNS changes cue-induced rating) versus the null hypothesis (Quintana & Williams, 2018). Bayesian testing combines a prior distribution of the expected effect with a measured data distribution (“likelihood”). As a result, we obtained a posterior distribution that integrates the prior and the observed data. Depending on the specification of the prior, a Bayes factor (BF) can be calculated. BF_10_ is defined as the ratio of the marginal likelihoods of the alternative hypothesis compared to the null hypothesis. Thus, a BF_10_ greater than one favors the alternative hypothesis whereas a BF_10_ lower than one favors the null hypothesis. Logically, the more extreme the BF is, the more conclusive is the evidence regarding the hypotheses that were defined a priori and BFs ~ 1 indicate that more data is necessary to provide conclusive evidence (Quintana & Williams, 2018). We used two-sided tests for all effects of interest. Additionally, to test the directed hypothesis that food wanting would be decreased during active taVNS, we used a one-sided t-test.

#### Statistical threshold and software

We collected (using Psychtoolbox v3, (Kleiner et al., 2007)) and preprocessed data in MATLAB v2018a. We conducted statistical tests and plotted results with JASP v0.09 – v.0.11(JASP team, 2019) and R v3.6.0 (R Core Team, 2019). As predefined by JASP, we set the prior of the Cauchy scale parameter to 0.707. To avoid that the inference is strongly dependent on the scaling parameter, we ran a prior robustness check as well, testing plausible narrower and wider prior settings.

## Results

### No effect of taVNS on ratings of food liking and wanting

To estimate taVNS-induced changes in food craving, we measured cue-induced liking and wanting ratings of food and non-food items. We calculated mean ratings during taVNS and sham stimulation for each participant for food and non-food categories to test for main effects of the stimulation. Neither food, nor non-food liking and wanting ratings showed significant differences between taVNS and sham conditions providing mostly moderate support for the null hypothesis (*p*s > .11, BF_10_ < .41; Table 1; Figure 2; for the Bayes factor robustness check see Figure S1 & S2). Moreover, differences between food and non-food ratings were not significantly altered by taVNS compared to sham providing moderate support against food-specific changes induced by taVNS (*p*s > .33, BF_10_ < .19). Notably, the data provide strong support against the alternative directed hypothesis that taVNS would decrease food wanting (BF_10_ = 0.05; Figure S3). Estimated effect sizes were low (dz < .18) suggesting limited practical relevance of any potential effect induced by acute taVNS.

**Table 1:** No main effect of taVNS on cue-induced liking and wanting ratings

| taVNS-induced changes<br>(taVNS – sham) | t | df | p | Mean<br>Difference | Lower<br>95% CI | Upper<br>95% CI | $d_z$ | $BF_{10}$ |
| --- | --- | --- | --- | --- | --- | --- | --- | --- |
| Food liking | 0.657 | 81 | .513 | 0.98 | -2.00 | 3.97 | 0.073 | 0.15 |
| Food wanting | 1.601 | 81 | .113 | 1.75 | -0.43 | 3.93 | 0.177 | 0.41 |
| Non-food liking | 1.501 | 81 | .137 | 1.95 | -0.64 | 4.53 | 0.166 | 0.36 |
| Non-food wanting | 0.211 | 81 | .833 | 0.26 | -2.18 | 2.69 | 0.023 | 0.12 |
| Food versus non-food<br>liking differences | -0.615 | 81 | .540 | -0.96 | -4.08 | 2.15 | -0.068 | 0.15 |
| Food versus non-food<br>wanting differences | 0.979 | 81 | .330 | 1.49 | -1.54 | 4.52 | 0.108 | 0.19 |

*Note.* Results of a Student's one sample t-test as well as Bayesian inference evaluating the mean differences between taVNS and sham stimulation and Bayes factor ( $BF_{10}$ ) for food liking, food wanting, non-food liking and non-food wanting as well as food versus non-food wanting and liking differences.

**Figure 2:**
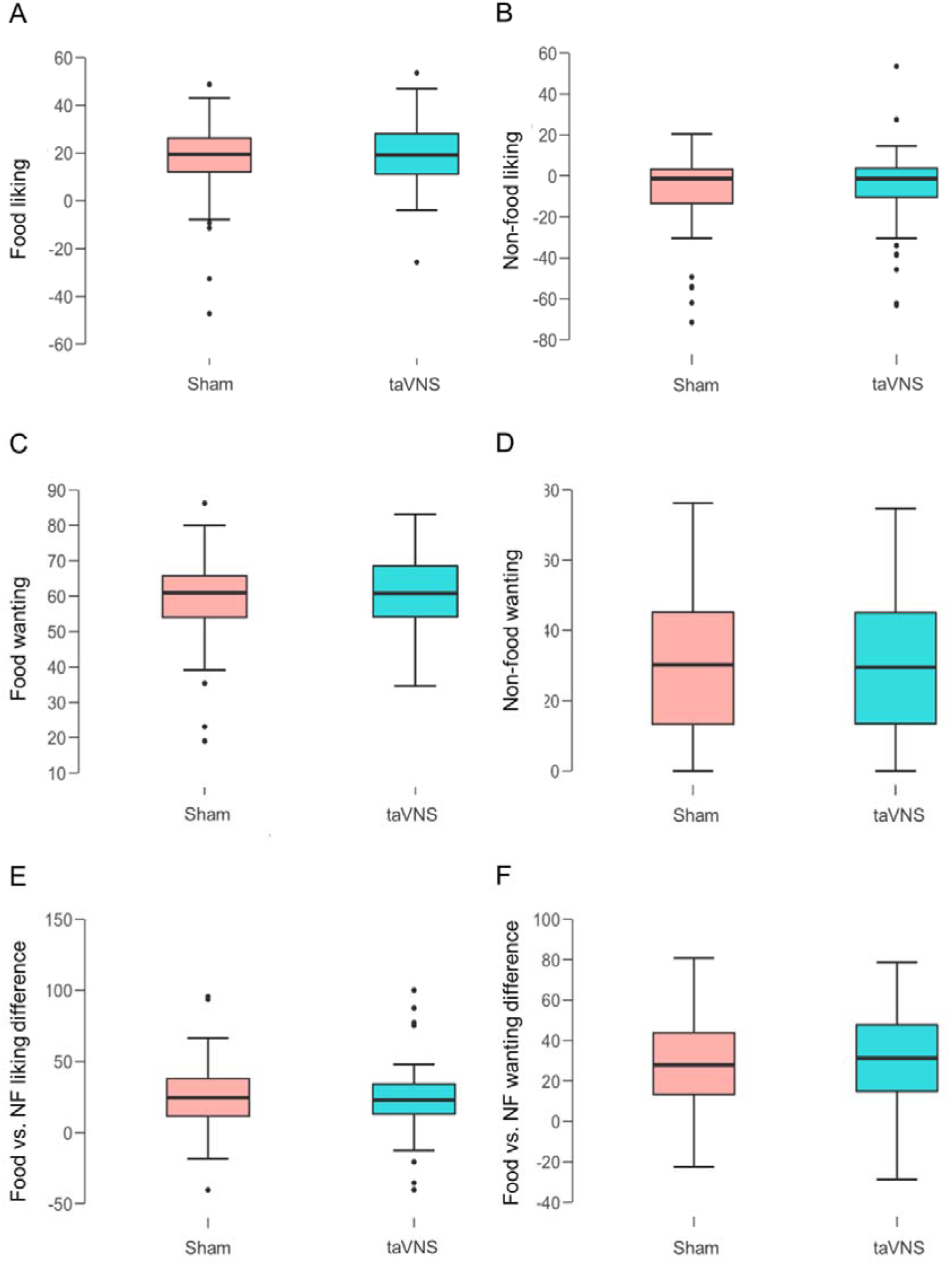
No effect of taVNS on cue-induced liking and wanting ratings. Boxplots depict individual averages of food and non-food liking (A & B) and wanting (C & D) ratings as well as differences between food and non-food (NF) liking (E) and wanting (F) during taVNS and sham stimulation.

### No effect of taVNS on liking and wanting ratings in post-hoc subgroup analyses

To rule out that the absence of taVNS main effects is due to condition-specific effects, we split the collected ratings according to stimulation side and sex (between-subject factors) as well as caloric density, perceived healthiness, and sweet versus savory foods (within-subject factors) and calculated each participants’ average ratings during taVNS and sham conditions again (for an overview on all subgroup analyses, see Figure 3, Table S1). Using these within- and between-subject factors, we found only one significant effect for liking ratings at an uncorrected *p*-threshold for non-food items during taVNS at the right side (*p* = .026). Likewise, for wanting ratings, we found 4 significant effects at an uncorrected *p*-threshold. We observed the lowest *p*-value for high-caloric food during taVNS at the right side (*p* = .016). Still, evidence in support of a taVNS effect was merely at an anecdotal level (BF_10_ = 2.68; Figure S4) suggesting that more data would be needed to provide sufficient evidence for the presence or absence of a condition-specific effect. To summarize, subgroup analyses provided anecdotal support for taVNS-induced changes at best. Thereby, our results provide mostly conclusive evidence for the absence of acute taVNS-induced effects on ratings of wanting and liking even within more narrowly defined conditions.

**Figure 3:**
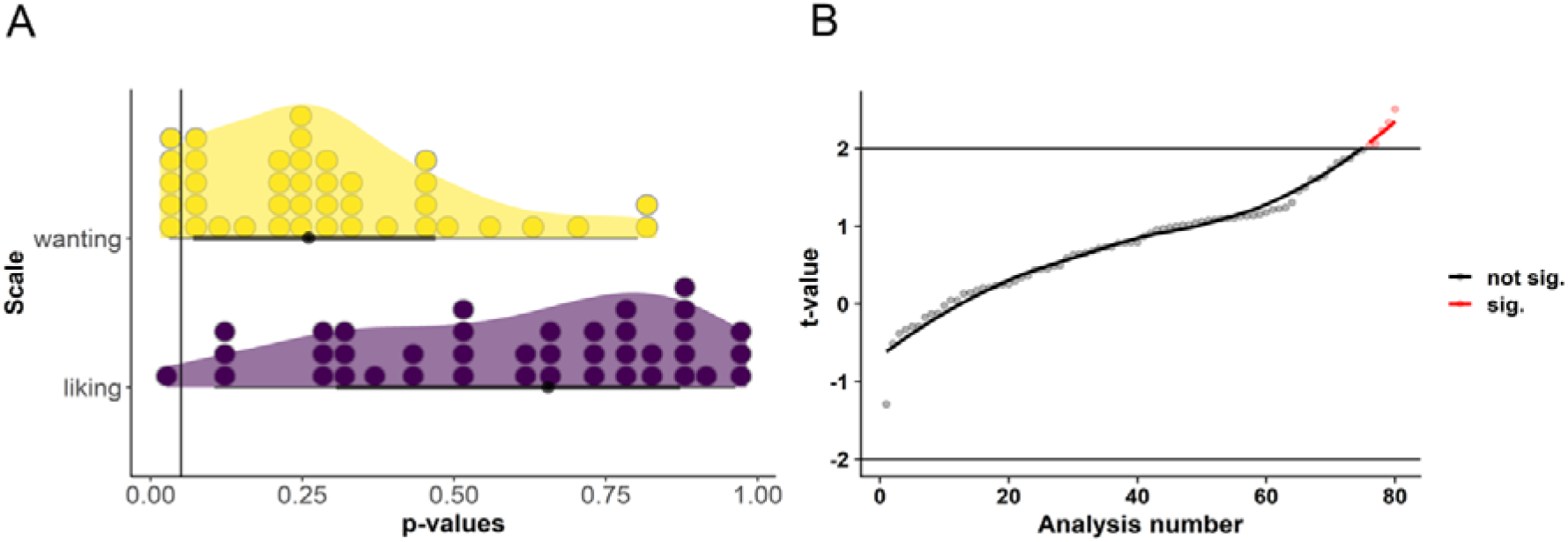
Post-hoc subgroup analyses provide mostly conclusive support for the null hypothesis that taVNS does not acutely effect ratings of wanting and liking. A: Distribution of *p*-values values for a stimulation main effect on ratings of liking and wanting within different subgroups defined by between-subject and within-subject factors. B: Corresponding distribution of t-values. The line denotes the uncorrected p-threshold for the 80 separate analyses. No result exceeded p-thresholds corrected for multiple comparisons or provided more than anecdotal evidence from Bayesian analyses for the presence of taVNS-induced changes.

Due to previously reported associations between BMI and sensitivity to homeostatic feedback signals (Schwartz & Porte, 2005), we reasoned that body composition may modulate the effects of taVNS on food picture wanting and liking. Again, using Bayesian inference, we observed no significant association between BMI and taVNS-induced changes in liking ratings (*r* = −0.111, BF_10_ = 0.22) or wanting ratings for food (*r* = −0.074 with a BF_10_ = 0.17; Figure 4). Moreover, BMI was not associated (−0.084 ≤ *r* ≤ 0.259, 0.14 ≤ BF_10_ ≤ 2.09) with ratings of liking or wanting in any of the other conditions we used for the subgroup analyses (stimulation side and sex as between-subject factors; caloric density, perceived healthiness, and sweet versus savory flavor of the food as within-subject factors). Thus, there was no indication that BMI modulates taVNS-induced changes in liking and wanting ratings of food.

**Figure 4:**
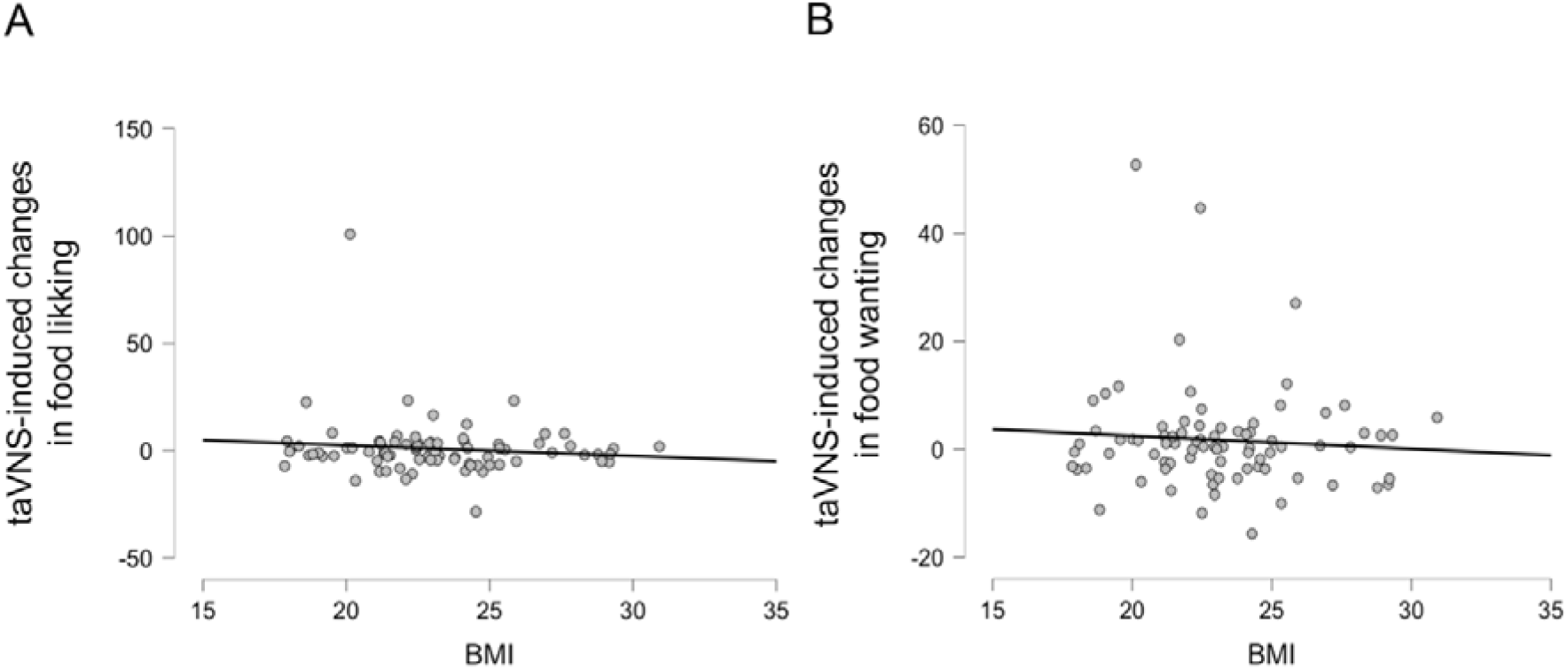
No association between BMI and taVNS-induced effects on ratings of liking and wanting for food. A: Correlation between BMI and taVNS-induced changes in liking ratings of food pictures. B: Correlation between BMI and taVNS-induced changes in wanting ratings of food pictures.

## Discussion

Vagal afferent signals play a vital role in the regulation of eating behavior by forwarding interoceptive feedback to tune goal-directed behavior. To better understand the contribution of vagal afferent activation in regulating wanting and liking of food, we investigated the impact of taVNS on cue-induced craving. As hypothesized, taVNS did not alter ratings of liking for food or ratings of liking and wanting for non-food items. However, in contrast to previous reports, wanting ratings for food were also not affected by acute taVNS, which is in accordance with an absence of taVNS effects on wanting ratings in an effort allocation task within the same study (Neuser et al., 2020). We conclude that effects on conscious ratings of liking and wanting elicited by acute taVNS are unlikely to account for previously reported weight loss due to chronic invasive VNS (Abubakr & Wambacq, 2008; Burneo et al., 2002; Pardo et al., 2007). These results call for further research on subacute and subconscious motivational effects of taVNS (Kühnel et al., 2020; Neuser et al., 2020) that may help elucidate the potential of taVNS for future eating-related interventions.

To date, our study is the largest effort to estimate taVNS effects on rated wanting and liking of food in a well-controlled experimental design (i.e., single-blind cross-over study). Still, our results provide mostly moderate evidence for the null hypothesis that conscious ratings of food are not acutely affected by taVNS. Moreover, since we observed non-significant increases in wanting and liking, our data provides strong evidence against the directed hypothesis that taVNS may acutely decrease food craving in healthy participants. The absence of taVNS-induced changes in rated liking and wanting was also supported by our post hoc subgroup analyses that evaluated potential condition-specific effects or potential associations with BMI and WtHR. These findings are well in line with recent smaller studies that reported no changes in food consumption or wanting ratings during acute taVNS in healthy participants (Alicart et al., 2020; Obst et al., 2020). Notably, the absence of taVNS-induced changes in ratings of cue-induced wanting and liking is also in line with the previously reported absence of taVNS effects on wanting in an effort allocation task despite a taVNS-induced increase in the invigoration to work for rewards at stake (Neuser et al., 2020). Collectively, these findings provide further support for the theorized subconscious role of reward signals originating from the gut (de Araujo et al., 2019).

Our observation that taVNS does not acutely decrease food craving appears to be at odds with previous reports of reduced food intake and body weight after VNS (Abubakr & Wambacq, 2008; Burneo et al., 2002; Pardo et al., 2007). However, this discrepancy could be due to differences between chronic stimulation regimes and the acute stimulation used in our work. For example, Vijgen et al. (2013) showed that chronic VNS increased energy expenditure in humans. Likewise, Li et al. (2015) found that chronic taVNS led to decreased body weight due to alterations in energy expenditure in obese rats. In contrast to this, we observed no effect of acute taVNS on energy expenditure but found a reduction in gastric myoelectric frequency (Teckentrup et al., 2020). This change in gastric frequency may not be instantaneously reflected in ratings but could conceivably alter eating and expected satiety (Janssen et al., 2011). Since other vagally-mediated mechanisms contribute to weight loss, slower food intake (Roslin & Kurian, 2001) and stronger subjective satiation after consumption of food (Pardo et al., 2007) could contribute to practically relevant effects of (ta)VNS on body weight. Furthermore, increased willingness to work for rewards during taVNS (Neuser et al., 2020) may translate to greater vigor in initiating physical activity, thereby providing a promising additional mode of action of taVNS to improve the control of body weight. Taken together, it seems unlikely that acute taVNS has a practically relevant effect on conscious evaluation of food.

Nevertheless, despite its notable strengths, our study is not without several limitations. First, we investigated taVNS-induced changes in food craving in healthy participants with a limited range in BMI. Future studies should include participants who are suffering from alterations in eating behavior related to mental or metabolic disorders as well to evaluate whether taVNS could improve pathological symptoms of eating. Second, we investigated taVNS-induced changes in cue-induced craving during acute stimulation. This is in contrast to most previous studies that reported anorexigenic effects in humans after (chronic) VNS (Abubakr & Wambacq, 2008; Burneo et al., 2002; Pardo et al., 2007). It is important to note that these studies investigated patients who suffered from either drug-resistant epilepsy (Abubakr & Wambacq, 2008; Burneo et al., 2002) or depression (Pardo et al., 2007) and did not include experimental control conditions. Still, Roslin and Kurian (2001) also observed that in mongrel dogs, chronic, but not acute VNS led to reduced food intake and a loss of body weight. Consequently, either chronic stimulation or longer periods of continuous taVNS may be necessary to elicit reliable effects on ratings of food and/or food intake.

In sum, we showed that taVNS does not acutely decrease subjective ratings of liking and wanting of food in healthy participants. Given a comparatively large sample testing within-subject effects elicited by taVNS, we provided mostly moderate evidence against an acute modulatory effect of taVNS. However, this does not preclude the potential presence of small acute effects for more specific categories of food such as high-caloric food, where we observed anecdotal evidence during taVNS applied at the right ear. Still, given the reported moderate-to-large effects of chronic invasive VNS on body weight in animals and humans, more mechanistic research is necessary to unravel subacute or subconscious effects of taVNS that could be used to improve future treatments of pathological alterations in eating behavior and food choice. In general, our results support the idea that vagal afferent activation elicits unconscious effects on food choice, which is in line with the theorized role of the “low road” to food choice (de Araujo et al., 2019).

## Acknowledgement

We thank Monja P. Neuser, Caroline Burrasch, Sandra Neubert, Moritz Herkner, and Leonie Osthof for help with data acquisition as well as Vincent Wolf for feedback on an earlier draft. The study was supported by the University of Tübingen, Faculty of Medicine fortune grant #2453-0-0. NBK received support from the Daimler and Benz Foundation, grant 32-04/19, the Else Kröner-Fresenius Stiftung, grant 2017_A67. MF received salary support from the University of Tübingen, Faculty of Medicine, IZKF Promotionsstipendium.

## Author contributions

NBK was responsible for the study concept and design. FKM & MF collected data under supervision by NBK. NBK conceived the method. FKM processed the data and performed the data analysis with contributions by AK under supervision by NBK. FKM & NBK wrote the manuscript. All authors contributed to the interpretation of findings, provided critical revision of the manuscript for important intellectual content and approved the final version for publication.

