## Supplementary Information for "Acute vagus nerve stimulation does not affect liking or wanting ratings of food in healthy participants"

**Supporting Information**

Franziska K. Müller^1^, Vanessa Teckentrup^1^, Anne Kühnel^1,2^, Magdalena Ferstl^1^,

& Nils B. Kroemer^1*^

^1^	Department of Psychiatry and Psychotherapy, University of Tübingen, Tübingen, Germany

^2^	Max Planck Institute of Psychiatry and International Max Planck Research School for Translational Psychiatry (IMPRS-TP), Munich, Germany

**Corresponding author***

Dr. Nils B. Kroemer,

Calwerstr. 14, 72076 Tübingen, Germany

### **Additional figures and analyses**

| A  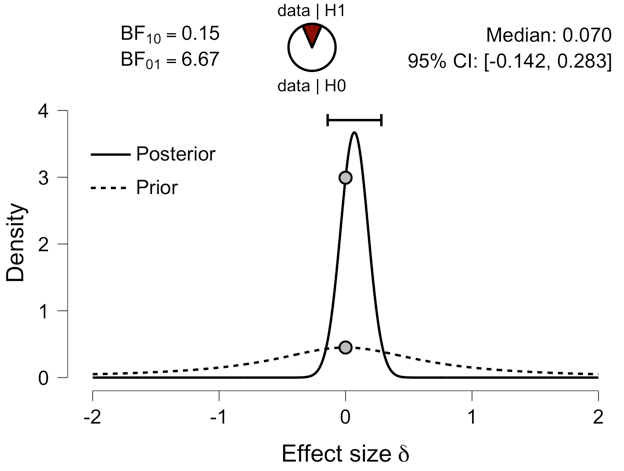 | B  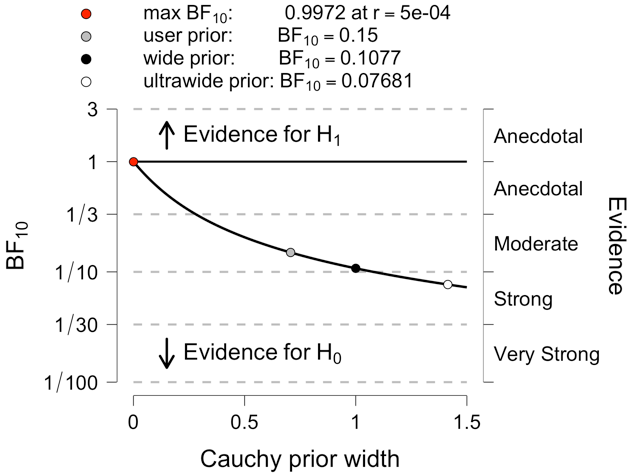 |
| --- | --- |

**Figure S1: Moderate evidence against taVNS effect on food liking**
Prior and posterior distribution of effects sizes of food liking rating differences during taVNS compared to sham stimulation (A). The Bayes factor robustness check indicates moderate to strong evidence that there is no effect of taVNS on food liking (B).

| A  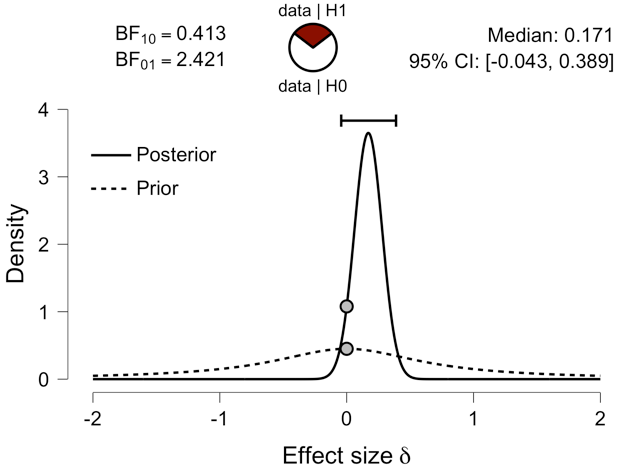 | B  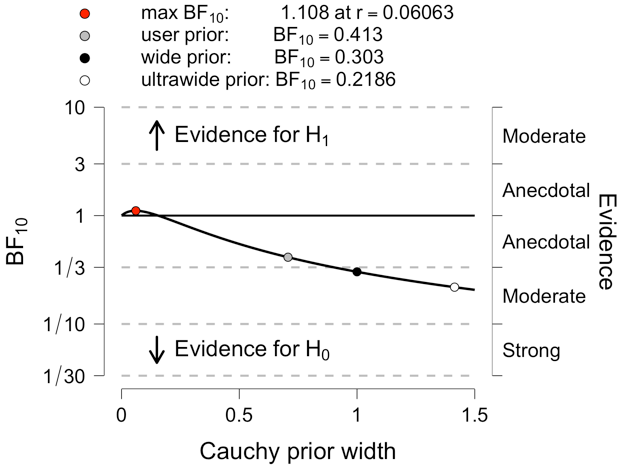 |
| --- | --- |

**Figure S2:** **Anecdotal evidence against an acute effect of taVNS on food wanting.**
Prior and posterior distribution of effect sizes of food wanting ratings differences during taVNS compared to sham stimulation (A). The Bayes factor robustness check indicates anecdotal to moderate support against an acute effect of taVNS on food wanting (B).

| A  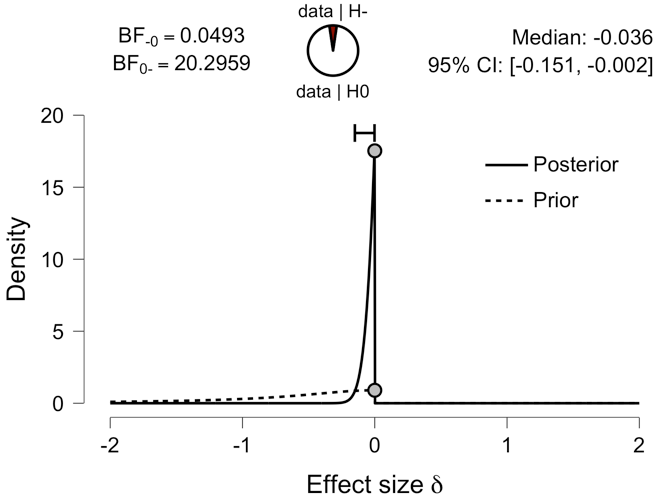 | B  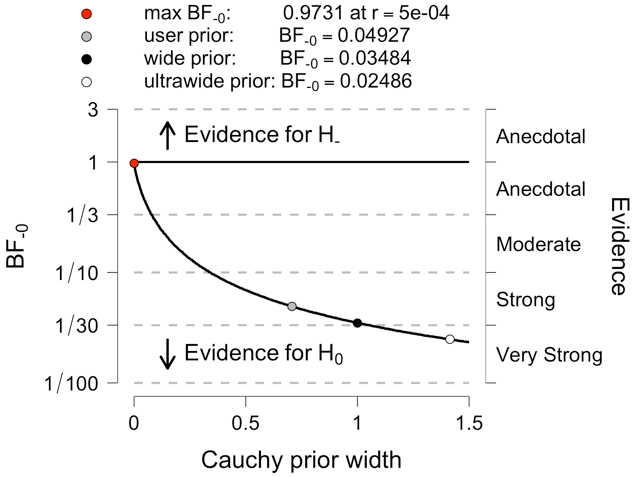 |
| --- | --- |

**Figure S3: Strong evidence that taVNS does not decrease food wanting**
Prior and posterior distribution of effect sizes of food wanting ratings during taVNS compared to sham stimulation considering a directional alternative hypothesis that taVNS decreases wanting ratings (A) and Bayes factor robustness check (B).

| A  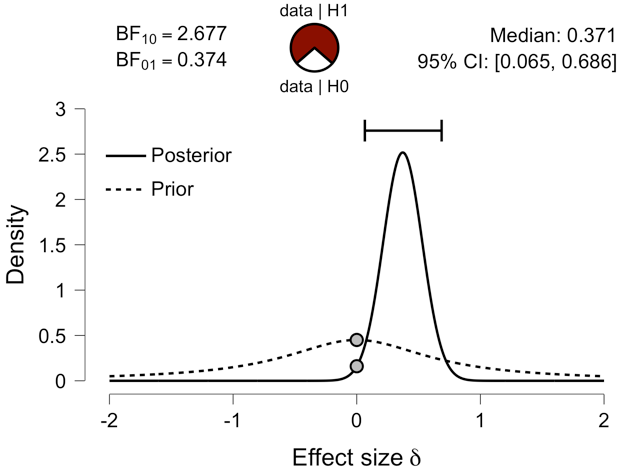 | B  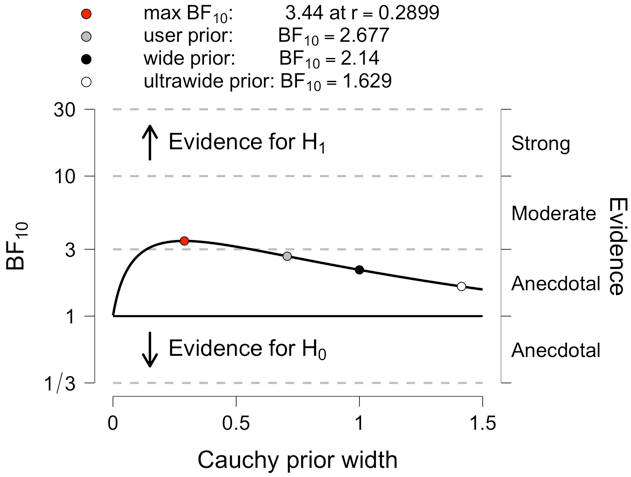 |
| --- | --- |

**Figure S4:** **Anecdotal evidence that right-side taVNS affects high caloric food wanting**
Prior and posterior distribution of effect sizes of High caloric food wanting differences between taVNS and sham stimulation at the right ear (A). The Bayes factor robustness check indicates anecdotal evidence in support of a stimulation effect of right side taVNS on high caloric food wanting (B).

**Table S1: Subgroup analyses of taVNS effects**

Results of one-sample t-tests testing differences between taVNS and sham stimulation for liking and wanting in subsets of the data based on sex, stimulation side, and stimulus characteristics. Only 6 of the 82 comparisons were nominally significant.

| **taVNS-induced changes (taVNS – sham)** | | | | | | | | | | |  |
| --- | --- | --- | --- | --- | --- | --- | --- | --- | --- | --- | --- |
| **Rating** | **Stimulus** | **StimSide** | **Sex** | **Caloric content** | **Perceived healthiness** | **Flavor** | **N** | | **Mean Diff** | | **p** |
| Wanting | Food | All | All | All | All | All | 82 |  | | 0.113 | |
| Wanting | Non-Food | All | All | - | - | - | 82 |  | | 0.833 | |
| Wanting | Food | All | All | High caloric | - | - | 82 | 2.075 | | 0.051 | |
| Wanting | Food | All | All | Low caloric | - | - | 82 | 1.464 | | 0.262 | |
| Wanting | Food | All | All | - | Healthy | - | 82 | 1.596 | | 0.195 | |
| Wanting | Food | All | All | - | Unhealthy | - | 82 | 1.983 | | 0.073 | |
| Wanting | Food | All | All | - | - | Sweet | 82 | 1.389 | | 0.259 | |
| **Wanting** | **Food** | **All** | **All** | **-** | **-** | **Savory** | **82** | **2.304** | | **0.045*** | |
| Wanting | Food | All | Female | All | All | All | 46 | 2.077 | | 0.245 | |
| Wanting | Non-Food | All | Female | - | - | - | 46 | -0.617 | | 0.704 | |
| Wanting | Food | All | Female | High caloric | - | - | 46 | 2.055 | | 0.221 | |
| Wanting | Food | All | Female | Low caloric | - | - | 46 | 2.156 | | 0.315 | |
| Wanting | Food | All | Female | - | Healthy | - | 46 | 2.189 | | 0.276 | |
| Wanting | Food | All | Female | - | Unhealthy | - | 46 | 2.021 | | 0.255 | |
| Wanting | Food | All | Female | - | - | Sweet | 46 | 2.247 | | 0.264 | |
| Wanting | Food | All | Female | - | - | Savory | 46 | 2.132 | | 0.231 | |
| Wanting | Food | All | Male | All | All | All | 36 | 1.332 | | 0.229 | |
| Wanting | Non-Food | All | Male | - | - | - | 36 | 1.377 | | 0.469 | |
| Wanting | Food | All | Male | High caloric | - | - | 36 | 2.101 | | 0.069 | |
| Wanting | Food | All | Male | Low caloric | - | - | 36 | 0.581 | | 0.63 | |
| Wanting | Food | All | Male | - | Healthy | - | 36 | 0.840 | | 0.481 | |
| **Wanting** | **Food** | **All** | **Male** | **-** | **Unhealthy** | **-** | **36** | **1.934** | | **0.09*** | |
| Wanting | Food | All | Male | - | - | Sweet | 36 | 0.292 | | 0.802 | |
| Wanting | Food | All | Male | - | - | Savory | 36 | 2.523 | | 0.059 | |
| Wanting | Food | Right | All | All | All | All | 40 | 1.822 | | 0.071 | |
| Wanting | Non-Food | Right | All | - | - | - | 40 | 2.660 | | 0.155 | |
| **Wanting** | **Food** | **Right** | **All** | **High caloric** | **-** | **-** | **40** | **2.822** | | **0.016*** | |
| Wanting | Food | Right | All | Low caloric | - | - | 40 | 0.623 | | 0.559 | |
| Wanting | Food | Right | All | - | Healthy | - | 40 | 1.091 | | 0.304 | |
| **Wanting** | **Food** | **Right** | **All** | **-** | **Unhealthy** | **-** | **40** | **2.546** | | **0.031*** | |
| Wanting | Food | Right | All | - | - | Sweet | 40 | 1.198 | | 0.289 | |
| **Wanting** | **Food** | **Right** | **All** | **-** | **-** | **Savory** | **40** | **2.603** | | **0.046*** | |
| Wanting | Food | Left | All | All | All | All | 42 | 1.682 | | 0.389 | |
| Wanting | Non-Food | Left | All | - | - | - | 42 | -2.029 | | 0.204 | |
| Wanting | Food | Left | All | High caloric | - | - | 42 | 1.364 | | 0.439 | |
| Wanting | Food | Left | All | Low caloric | - | - | 42 | 2.266 | | 0.337 | |
| Wanting | Food | Left | All | - | Healthy | - | 42 | 2.077 | | 0.347 | |
| Wanting | Food | Left | All | - | Unhealthy | - | 42 | 1.447 | | 0.437 | |
| Wanting | Food | Left | All | - | - | Sweet | 42 | 1.570 | | 0.47 | |
| Wanting | Food | Left | All | - | - | Savory | 42 | 2.019 | | 0.286 | |
| Liking | Food | All | All | All | All | All | 82 |  | | 0.513 | |
| Liking | Non-Food | All | All | - | - | - | 82 |  | | 0.137 | |
| Liking | Food | All | All | High caloric | - | - | 82 | 0.374 | | 0.798 | |
| Liking | Food | All | All | Low caloric | - | - | 82 | 1.654 | | 0.327 | |
| Liking | Food | All | All | - | Healthy | - | 82 | 1.652 | | 0.322 | |
| Liking | Food | All | All | - | Unhealthy | - | 82 | 0.194 | | 0.896 | |
| Liking | Food | All | All | - | - | Sweet | 82 | 1.745 | | 0.312 | |
| Liking | Food | All | All | - | - | Savory | 82 | 0.195 | | 0.887 | |
| Liking | Food | All | Female | All | All | All | 46 | 1.907 | | 0.432 | |
| Liking | Non-Food | All | Female | - | - | - | 46 | -0.278 | | 0.864 | |
| Liking | Food | All | Female | High caloric | - | - | 46 | 1.026 | | 0.657 | |
| Liking | Food | All | Female | Low caloric | - | - | 46 | 2.891 | | 0.295 | |
| Liking | Food | All | Female | - | Healthy | - | 46 | 2.973 | | 0.275 | |
| Liking | Food | All | Female | - | Unhealthy | - | 46 | 0.700 | | 0.768 | |
| Liking | Food | All | Female | - | - | Sweet | 46 | 3.050 | | 0.279 | |
| Liking | Food | All | Female | - | - | Savory | 46 | 0.762 | | 0.719 | |
| Liking | Food | All | Male | All | All | All | 36 | -0.195 | | 0.897 | |
| **Liking** | **Non-Food** | **All** | **Male** | **-** | **-** | **-** | **36** | **4.792** | | **0.026*** | |
| Liking | Food | All | Male | High caloric | - | - | 36 | -0.459 | | 0.775 | |
| Liking | Food | All | Male | Low caloric | - | - | 36 | 0.074 | | 0.963 | |
| Liking | Food | All | Male | - | Healthy | - | 36 | -0.036 | | 0.982 | |
| Liking | Food | All | Male | - | Unhealthy | - | 36 | -0.453 | | 0.771 | |
| Liking | Food | All | Male | - | - | Sweet | 36 | 0.077 | | 0.963 | |
| Liking | Food | All | Male | - | - | Savory | 36 | -0.529 | | 0.742 | |
| Liking | Food | Right | All | All | All | All | 40 |  | | 0.523 | |
| Liking | Non-Food | Right | All | - | - | - | 40 | 3.451 | | 0.114 | |
| Liking | Food | Right | All | High caloric | - | - | 40 | 0.291 | | 0.815 | |
| Liking | Food | Right | All | Low caloric | - | - | 40 | 0.892 | | 0.53 | |
| Liking | Food | Right | All | - | Healthy | - | 40 | 1.262 | | 0.369 | |
| Liking | Food | Right | All | - | Unhealthy | - | 40 | -0.168 | | 0.903 | |
| Liking | Food | Right | All | - | - | Sweet | 40 | 1.916 | | 0.107 | |
| Liking | Food | Right | All | - | - | Savory | 40 | -0.681 | | 0.613 | |
| Liking | Food | Left | All | All | All | All | 42 | 1.243 | | 0.653 | |
| Liking | Non-Food | Left | All | - | - | - | 42 | 0.516 | | 0.734 | |
| Liking | Food | Left | All | High caloric | - | - | 42 | 0.453 | | 0.863 | |
| Liking | Food | Left | All | Low caloric | - | - | 42 | 2.380 | | 0.433 | |
| Liking | Food | Left | All | High caloric | - | - | 42 | 0.453 | | 0.863 | |
| Liking | Food | Left | All | Low caloric | - | - | 42 | 2.380 | | 0.433 | |
| Liking | Food | Left | All | - | Healthy | - | 42 | 2.024 | | 0.50 | |
| Liking | Food | Left | All | - | Unhealthy | - | 42 | 0.538 | | 0.837 | |
| Liking | Food | Left | All | - | - | Sweet | 42 | 1.582 | | 0.622 | |
| Liking | Food | Left | All | - | - | Savory | 42 | 1.030 | | 0.664 | |
